# Sonic hedgehog medulloblastoma cells in co-culture with cerebellar organoids converge towards *in vivo* malignant cell states

**DOI:** 10.1101/2024.04.01.587603

**Authors:** Max J. van Essen, Alina Nicheperovich, Benjamin Schuster-Böckler, Esther B. E. Becker, John Jacob

## Abstract

**Background:** In the malignant brain tumour sonic hedgehog medulloblastoma (SHH-MB) the properties of cancer cells are influenced by their microenvironment, but the nature of those effects and the phenotypic consequences for the tumour are poorly understood. The aim of this study was to identify phenotypic properties of SHH-MB cells that were driven by the non-malignant tumour microenvironment.

**Methods:** Human induced pluripotent cells (iPSC) were differentiated to cerebellar organoids to simulate the non-malignant tumour microenvironment. Tumour spheroids were generated from two distinct, long-established SHH-MB cell lines which were co-cultured with cerebellar organoids. We profiled the cellular transcriptomes of malignant and non-malignant cells by performing droplet-based single-cell RNA-sequencing (scRNA-seq). The transcriptional profiles of tumour cells in co-culture were compared with those of malignant cells cultured in isolation and with public SHH-MB datasets of patient tumours and patient-derived xenograft (PDX) models.

**Results:** SHH-MB cell lines in organoid co-culture adopted patient tumour-associated phenotypes and showed increased heterogeneity compared to monocultures. Sub-populations of co-cultured SHH-MB cells activated a key marker of differentiating granule cells, *NEUROD1* that was not observed in tumour monocultures. Other sub-populations expressed transcriptional determinants consistent with a cancer stem cell (CSC)-like state that resembled cell states identified *in vivo*.

**Conclusion:** For SHH-MB cell lines in co-culture, there was a convergence of malignant cell states towards patterns of heterogeneity in patient tumours and PDX models implying these states were non-cell autonomously induced by the microenvironment. Therefore, we have generated an advanced, novel *in vitro* model of SHH-MB with potential translational applications.

## Introduction

Medulloblastoma (MB), a heterogeneous cerebellar tumour consisting of four major subtypes is the commonest malignant brain tumour of childhood, ^1^ of which nearly a third are classed as Sonic Hedgehog medulloblastoma (SHH-MB).^2^ During development, SHH acting as a mitogen promotes massive expansion of the upper rhombic lip-derived cerebellar granule cell population in the external granule layer (EGL).^3^ Somatic or germ-line mutations and focal somatic copy number alterations in SHH pathway tumour suppressors or oncogenes are the commonest events leading to aberrant SHH pathway activation and tumorigenesis in granule progenitors.^2^ High-risk variants of SHH-MB still carry a dismal prognosis, which emphasises the importance of gaining a greater understanding of how tumours progress so that improved therapies can be developed.

In patients and mouse models, SHH-MB contains SOX2^+^ tumour-propagating neural stem-like cells, equivalent to cancer stem cells (CSC) and more differentiated cells.^2^ By hijacking mechanisms of self-renewal in neural stem cells (NSC) and pluripotent stem cells, CSC can replenish tumours in transplantation assays.^4^ During mammalian neurogenesis, initial ubiquitous SOX2 expression in the early embryonic central nervous system attenuates as the balance shifts from self-renewal to differentiation.^5^ By contrast, in SHH-MB persistent SOX2-expressing granule progenitor-derived CSC can reconstitute tumours after therapy and support a hierarchical model of tumorigenesis.^6^ Furthermore, cross-talk between the microenvironment and other brain tumour types suggests that cell extrinsic factors could also influence malignant cell states.^4^ However, the complexity of the *in vivo* tumour microenvironment poses a challenge in discerning the involvement of distinct non-malignant components.

To address how the microenvironment affects tumour progression, we sought to evaluate the phenotypes of SHH-MB cells using single-cell transcriptomics, under defined conditions that we anticipated might simulate *in vivo* tumour growth. To this end we leveraged cerebellar organoid differentiation from human induced pluripotent stem cells (hiPSC) through their co-culture with two SHH-MB cell lines ^7^ that were compared with conventional *in vitro* tumour monocultures. Organoids can be grown in xeno-free culture medium and they contain defined neuronal and glial cell types. Although lacking immune cells or blood vessels, they provide more physiological conditions than conventional *in vitro* models enabling the evaluation of tumour cell phenotypes in a simulated native microenvironment.^8^

## Materials and Methods

### Organoid differentiation

Cerebellar organoid differentiation was performed as described previously.^9^ In brief, human iPSC line AH017-3 cells ^10^ were detached by incubation with pre-warmed TrypLE for 5 minutes. Cells were collected by centrifugation and resuspended in induction medium (50% IMDM, 50% F12, 7ug/mL Insulin, 5mg/mL BSA, 1X Chemically defined Lipid concentrate, 450uM Monothioglycerol, 15ug/mL Apo-transferrin, 1X Penicillin/Streptomycin) at a final concentration of 100,000 cells/ml. 10,000 cells per well were plated in an ultra-low-attachment V-bottom 96 well plate and returned to the incubator. After two days, FGF2 was added at a final concentration of 50 ng/ml. One-third medium change was performed at day 7 and full medium change at day 14. On this day, the organoids were also transferred into a low-attachment 48-well plate. On day 21, medium was changed to differentiation medium (Neurobasal + 1% N2 supplement + 1% GlutaMAX + 1X Penicillin/Streptomycin) and subsequent full medium changes were performed on days 28 and 35. After day 35, medium was changed twice/week.

### Medulloblastoma cell lines, tumour spheroids and organoid-tumour spheroid co-culture

DAOY-GFP cells (gift from Vincenzo D’Angiolella, DAOY cell line originally purchased from ATCC) were grown in monolayer culture in MEM, 1% GlutaMAX, 1X Penicillin/Streptomycin and 1% FBS (cell line authentication in Supplementary Table 1). ONS-76-Luciferase-GFP cells (gift from Louis Chesler; ONS-76 cell line originally purchased from JCRB Cell Bank by Steven Clifford) were grown in monolayer culture in RPMI + 1% GlutaMAX + 1X Penicillin/Streptomycin + 1% FBS (Supplementary Table 1). Medium was changed twice per week. To generate tumour spheroids, cells were detached by incubation with pre-warmed TrypLE for 5 minutes and collected by centrifugation. Cells were resuspended in differentiation medium at a concentration of 200,000 cells/ml and plated in an ultra-low-attachment V-bottom 96-well plate at 20,000 cells per well. In preparation for cerebellar organoid-tumour spheroid co-culture, one week prior to the start of co-culture, tumour spheroids were generated as described above. One tumour spheroid was placed with one day 35 organoid in a low-attachment 24-well plate in identical differentiation medium. The size ratio of the organoid to the tumour spheroid of approximately five-to-one ensured that the organoid microenvironment could encompass the tumour spheroid to facilitate their interaction. To allow fusion of the tumour spheroid and the organoid, the plate was incubated for two days at a 45-degree angle. Medium was changed twice per week and the cells were maintained in co-culture, alongside tumour spheroid and tumour monolayer cultures for a further 25 days (Fig. 1A).

**Figure 1.**
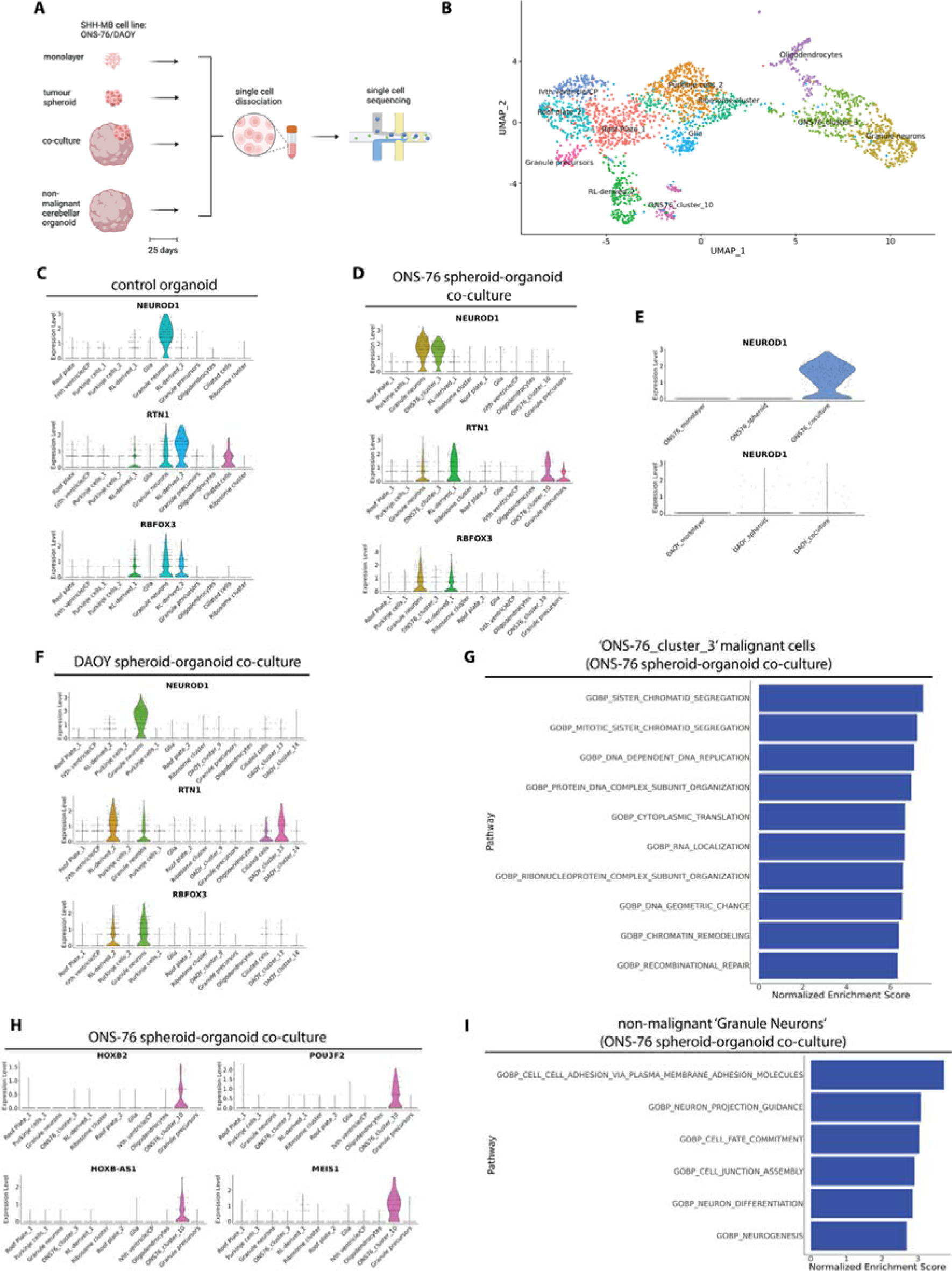
Upregulation of *NEUROD1* in ONS-76 cells in co-culture compared to monolayer or tumour spheroid growth conditions. (A) Schematic of experimental workflow. This figure was created with BioRender.com. (B) UMAP visualisation of single-cell transcriptomes of ONS-76 cells in co-culture identified two malignant cell clusters annotated as ‘ONS76_cluster_3’ and ‘ONS76_cluster_10’. (C) Violin plot showing co-expression of *NEUROD1*, *RTN1* and *RBFOX3* in the ‘Granule neurons’ cluster of the control organoid. (D) Violin plot of expression of *NEUROD1,* and absent *RTN1* and *RBFOX3* expression in ‘ONS76_cluster_3’ in the ONS-76 tumour spheroid-organoid co-culture (E) Violin plots of the expression of *NEUROD1* in ONS-76 cells (upper panel) and DAOY cells (lower panel) grown as monolayers, tumour spheroids or in tumour spheroid-organoid co-culture. (F) Expression of *NEUROD1* in the DAOY tumour spheroid-organoid co-culture sample is restricted to the non-malignant ‘Granule neurons’ cluster. (G) fgsea gene set enrichment of upregulated gene ontology (GO) biological pathway terms evaluated by normalised enrichment scores of the *NEUROD1*+ cluster of malignant cells (‘ONS76_cluster_3’) in the ONS-76 tumour spheroid-organoid co-culture sample. (H) Violin plot of marker genes of the second malignant cluster, ‘ONS76_cluster_10’ in the ONS-76 tumour spheroid-organoid co-culture sample. (I) fgsea gene set enrichment of upregulated GO biological pathway terms evaluated by normalised enrichment scores of non-malignant *NEUROD1*+ cells (‘Granule neurons’) in the ONS-76 tumour spheroid-organoid co-culture sample.

### Single cell isolation

Day 60 organoids and organoid-tumour co-culture samples were collected using a wide bore pipette tip and placed in a 6-well plate. Five tumour spheroid-organoid co-cultures for each cell line and a similar number of control organoids were each cut in four pieces and placed in an Eppendorf tube. After removal of excess medium, 200 μl of pre-warmed (to 37°C) Neuron isolation enzyme was added and the tube was placed in a 37°C water bath. Organoids were incubated with the enzyme for 30 minutes with intermittent agitation. The enzyme was removed and organoids were washed once with HHGN medium (HBSS + 2.5mM HEPES + 35mM Glucose + 4mM NaHCO_3_). Organoids were then dissociated in differentiation medium by gentle trituration. Afterwards, the cell suspension was enriched for viable cells using the dead cell removal kit (Miltenyi Biotec) following the manufacturers protocol. Cell viability was determined using Trypan Blue staining and counted using an automated cell counter (Applied Biosystems Countess 3). Cells were centrifuged at 300 x *g* for 5 minutes and resuspended in the appropriate volume of resuspension buffer (HBSS + 0.04% BSA) to a final concentration of 2000 cells/μl before proceeding with single-cell RNA sequencing. Thirty tumour spheroids were collected using a wide bore pipette tip and placed in an Eppendorf tube. Excess medium was removed and 200 μl of pre-warmed Neuron isolation enzyme (Gibco) was added. The tube was placed in a 37°C water bath for 20 minutes and agitated every 5 minutes. Subsequent processing was performed following the same steps as described above. DAOY and ONS-76 cell monolayer cultures were dissociated by incubation with pre-warmed TrypLE for 5 minutes. TrypLE was then diluted in PBS before centrifugation at 400 x *g* for 5 minutes to collect the cells. Cells were washed once in differentiation medium before processing as described above. Single cell barcoding and reverse transcription of poly-A mRNA was performed using the 10X Chromium controller (3’ version 3.1 chemistry) and sequencing of single-cell libraries was performed on an Illumina NovaSeq6000 sequencing system according to the manufacturer’s instructions.

### Data pre-processing and analysis

Initial sequencing data exploration was performed using Cell Ranger v6.1.1. For each dataset we removed cells with 10% or higher mitochondrial gene expression ^11^ and genes expressed in fewer than 10 cells. Sequencing confirmed the greater proportion of non-malignant to malignant cell types in co-culture which was expected given the relative sizes of the organoid and tumour spheroid (Supplementary Table 2). Downstream analysis largely implemented the workflow of Seurat v4, which is robust to limited cell numbers.^12^ We used the ‘sctransform’ modelling framework, which corrects counts for each gene in a cell by regressing out sequencing depth.^13^ The difference between G2M and S phase scores was regressed out so that differences in cell cycle phase among proliferating cells was eliminated, while at the same time retaining signals separating cycling and non-cycling cells. For differential gene expression testing between clusters a Bonferroni-corrected p-value < 0.05 was considered significant. To perform gene set enrichment analysis, we first used the R package PRESTO which performs a fast Wilcoxon rank sum test on the Seurat clusters.^14^ We then tested functional enrichment of Seurat clusters of interest using gene sets from MsigDB and the fgsea package.^15,16^ Visualisation of pathway enrichment analysis was performed using g:Profiler to search for Gene Ontology pathways that were significantly enriched in a gene list before visualising the results using EnrichmentMap in Cytoscape.^17,18^ Pathway gene sets with fewer than three genes and more than 350 genes were excluded in line with the default g:Profiler and Cytoscape workflows. For g:Profiler, a Benjamini-Hochberg false discovery rate adjusted p-value of < 0.05 was considered significant. Pseudotime analysis was conducted using Monocle 3.^19^ Dataset integrations were, in general, performed using standard workflows in Seurat v4, except when integrating malignant cell populations (see Fig. 5) where we observed strong blending of distinct malignant cell states. Instead, we used the fastMNN algorithm,^20^ which was found to preserve biological differences between cancer cell populations, while ensuring effective batch correction.^21^

### Statistical Analysis

Statistical significance testing was performed using R.

## Results

We co-cultured malignant tumour spheroids of ONS-76 cells or DAOY cells, which harbour wild type and mutant copies of *TP53*, respectively with non-malignant cerebellar organoids differentiated from human iPSC (Fig. 1A; Supplementary Fig. S1A). As controls, tumour cells were grown as monolayers or tumour spheroids while non-malignant organoids were grown separately. These long-established cell lines were chosen because of their evolutionary divergence from patient tumours. Our approach was designed to enhance the detection of malignant cell states induced by the non-malignant microenvironment, when comparing monocultures to the co-culture conditions. Using primary SHH-MB or PDX cultured cells could potentially mask such differences, given their closer resemblance to *in vivo* cell states, thus obscuring microenvironmental impacts on tumour cell properties. After 25 days in co-culture, cells were dissociated and processed for single-cell RNA sequencing (scRNA-seq) on the 10X Chromium platform. After filtering the dataset in accordance with our quality control thresholds we selected 13,534 cells for further analysis (Supplementary Fig. S1B – E; Supplementary Table 2). We used Seurat to visualise the relationship between samples by clustering and projected cells in a common UMAP space labelled according to their sample of origin (Supplementary Fig. S1F, G). ONS-76 cells and DAOY cells cultured as monolayers or tumour spheroids formed well-demarcated clusters. In contrast, there was extensive overlap of co-cultured cells with the non-malignant organoid control, which was anticipated given the large numbers of cells recovered belonging to organoid cellular subtypes. Only a handful of cells from the non-malignant control organoid clustered with the exclusively malignant samples, confirming that the clustering reflected biological differences between the samples. In the co-cultured samples, several times the number of cells clustered with the malignant samples suggesting the latter cells were malignant cells in co-culture.

### ONS-76 and DAOY cells cultured as monolayers or tumour spheroids were enriched for proliferative, protein biosynthetic and wound healing processes

We performed an integrated analysis of ONS-76 monolayer and tumour spheroid transcriptomes in Seurat ^13^ and compared the transcriptional programmes of malignant cells using functional enrichment analysis in the two conditions.^16^ ONS-76 cells grown as monolayers were enriched for Gene Ontology (GO) terms for cell cycle associated genes (Supplementary Fig. S2A and Supplementary Table 3). In comparison, genes upregulated in tumour spheroids were associated with altered cell morphology and wound healing for example *IFITM3, IGFBP2, IGFBP5, STC1* and *SPARC* (Supplementary Fig. S2B – C and Supplementary Table 3) consistent with an epithelial-to-mesenchymal transition (EMT) state and tumour stroma production.^22^ Upon clustering of the ONS-76 tumour spheroid sample (Supplementary Fig. S2D), expression of these genes was distributed over all clusters (Supplementary Fig. S2E) supporting relative homogeneity of malignant cell states with mostly arbitrary distinctions between clusters, with two exceptions. The first consisted of a cluster of cycling cells, denoted ‘cluster 4’ that was marked by the expression of genes including *MKI67*, *MYBL2*, *BIRC5* and *RRM2* consistent with a proliferative cell state (Supplementary Fig. S2D, G). Although the same genes were expressed in the ONS-76 monolayer sample, they were not restricted to a single cluster (Supplementary Fig. S2I). A second ONS-76 tumour spheroid cluster, ‘cluster 0’ expressed ribosomal genes and was enriched for protein synthesis pathways (Supplementary Fig. S2D and Supplementary Table 4), which is a feature common to many cancers.^23^ Less heterogeneity was evident in ONS-76 cells cultured as monolayers, which could be split into two clusters also distinguished by their relative enrichment for ribosomal genes and protein synthesis (Supplementary Fig. S2H; Supplementary Table 4). In comparison to ONS-76 tumour spheroids, expression of proliferation genes spanned both clusters, although the proliferation signal of individual cells was weaker (Supplementary Fig. S2H, lower panel).

Broadly similar findings were observed upon analysis of DAOY cells cultured as monolayers and tumour spheroids. An integrated analysis of the cellular transcriptomes revealed protein synthesis gene sets as the only significant enrichment (Supplementary Table 5) in DAOY tumour spheroids. In common with ONS-76 cells, wound healing gene sets were also upregulated, but in DAOY monolayers rather than in DAOY tumour spheroids. Furthermore, there was a degree of enrichment in oncogenic signature gene sets that was statistically non-significant for transcriptional responses of cerebellar granule progenitors to activation by SHH, which acts as a mitogen for these cells (Supplementary Table 5).

Clustering of the DAOY tumour spheroid sample also revealed enrichment of protein synthesis and cell cycle related pathways, respectively in two out of five DAOY tumour spheroid clusters (Supplementary Fig. S2F; Supplementary Table 5). Therefore, in tumour spheroids of both cell lines, those cells with a signature of enhanced proliferation were restricted to specific clusters in UMAP space, suggesting an association with a distinct cell state related to malignant cell proliferation. By contrast, DAOY monolayer cells that expressed a proliferative signature were spread diffusely across multiple clusters in common with ONS-76 monolayer cultured cells (Supplementary Fig. S2J). Although DAOY monolayer cells formed more clusters than ONS-76 cells (six in DAOY monolayers compared to two in the corresponding ONS-76 sample), only three out of six of the former clusters displayed significant gene set enrichment. Of these, two clusters were enriched for protein synthesis pathways and the remaining cluster was only significantly enriched for cell adhesion-related gene sets, which is a hallmark of cancer (Supplementary Table 5).^24^ Assessment of the maturity of malignant cells from tumour spheroid and monolayer samples of both cell lines revealed they expressed the granule cell markers, *MEIS1,*^25^ *MEIS2* ^26^ and *JKAMP* ^27^ (Supplementary Fig. S2K). However, *CNTN1* which marks differentiating granule cells ^28^ was selectively expressed only in ONS-76 samples, suggesting the maturation of DAOY cells was relatively impaired (Supplementary Fig. S2K).

### ONS-76 cells in co-culture, but not in monolayers or tumour spheroids expressed *NEUROD1*

To determine whether malignant cells grown in a non-malignant organoid microenvironment led to altered tumour cell phenotypes, we clustered the ONS-76 sample in co-culture and plotted the UMAP projection, comparing the clusters obtained with those in the non-malignant control organoid (Fig. 1B; Supplementary Fig. S3A). The manual annotations of non-malignant cells were highly similar to our earlier annotations of day 90 organoids.^29^ None of the clusters of malignant cells in co-culture were identified as having a match in the control organoid.

In the control organoid, granule neurons could be defined by their co-expression of *NEUROD1*, *RTN1* and *RBFOX3* (Fig. 1C).^27^ In the co-culture condition also, a cluster of cells co-expressing these genes could be identified consistent with non-malignant granule neurons (Fig. 1D). Additionally, a distinct cell cluster located adjacent to non-malignant granule neurons in UMAP space could be distinguished from the latter cells by their expression of *NEUROD1* only, which we labelled ‘ONS76_cluster_3’ (Fig. 1D; Supplementary Fig. S3B; Supplementary Table 6). Furthermore, upon computational integration of ONS-76 samples malignant *NEUROD1*^+^ cells were absent in ONS-76 monolayers and tumour spheroids (Fig. 1E) and only very few were present in the DAOY samples (Fig. 1E, F).

Although *NEUROD1* marks post-mitotic granule neurons in mouse,^30^ in humans this gene is also expressed in cycling granule progenitors.^31^ Gene set enrichment analysis of ‘ONS76_cluster_3’ revealed GO terms associated with cycling cells (Fig. 1G; Supplementary Fig. S3C). By contrast, the cluster annotated as ‘Granule neurons’ in the non-malignant organoid in co-culture and in the control organoid were enriched for terms associated with differentiated neurons (Fig. 1I; Supplementary Fig. S3E, F). To validate these findings, we computationally examined the cell cycle phase associated with *NEUROD1* expression in non-malignant and malignant cells.^13^ In the cluster annotated as ‘Granule neurons’ in the control organoid (Fig. 1C), the fraction of *NEUROD1*^+^ cells that were in G1 phase was 0.60, in G2/M phase the fraction was 0.17 and the S phase fraction was 0.23. However, in ONS-76 cells in co-culture the fraction of *NEUROD1*^+^ cells in ‘ONS76_cluster_3’ in G1 phase was smaller (0.38), and the fraction of cells in G2/M and S phases was higher (0.30 and 0.32, respectively; p = 5.925 x 10^-12^ Chi-squared test). Therefore, ONS-76 cells activated *NEUROD1,* a key molecular determinant of granule cell identity only when embedded in a more physiological microenvironment, but its expression was less tightly coupled to other cellular processes that imparted a differentiated neuronal identity. By comparison, analysis of patient SHH-MB scRNA-seq public datasets showed that they also expressed *NEUROD1* (Supplementary Fig. 4A-E) ^32,33^ and that for one of the datasets ^32^ most of these cells were in G1 phase (range 0.59 – 0.69). Consistent with maturation of malignant cells, patient tumours also expressed *RBFOX3* and *RTN1* (Supplementary Fig. S4A-E). Given the absence of *NEUROD1* upregulation in DAOY cells in co-culture, we analysed its expression in two *TP53* mutant SHH-MB in a second public MB scRNA-seq dataset to test for an association between presence of the mutation and *NEUROD1* expression *in vivo*.^33^ The tumours varied widely in their expression of *NEUROD1* displaying both elevated and minimal expression (Supplementary Fig. S4D). Therefore, absence of *NEUROD1* expression in *TP53* mutant DAOY cells in tumour spheroid-organoid co-culture was not inconsistent with *in vivo* findings.

A second, smaller cluster of ONS-76 tumour cells in co-culture, ‘ONS76_cluster_10’ was identified by differential upregulation of *HOXB2* and *POU3F2*, in common with multiple non-CNS cancers including leukaemia, lung and breast cancer (Fig. 1H; Supplementary Table 6).^34,35^ Cells in this cluster also expressed *MEIS1*, which is expressed by differentiating granule cells ^36^ and *HOXB-AS1*, which is a member of a large family of non-coding transcripts that are abnormally expressed in a wide range of cancers.^37^ Consistent with the expression of *MEIS1*, gene set enrichment analysis suggested this cluster contained mature cells (Supplementary Fig. S3G, H). Overall, subpopulations of malignant cells expressed *in vivo* markers of differentiation when they were surrounded by the more physiologic microenvironment of the organoid.

### A sub-population of DAOY cells in co-culture expressed multiple cancer stem cell-like markers

The analysis of the DAOY tumour spheroid-organoid co-culture sample resulted in clustering of DAOY tumour cells into three distinct groups in UMAP space designated as ‘DAOY_cluster_9’, ‘DAOY_cluster_13’, and ‘DAOY_cluster_14’ (Fig. 2A). Highly distinctive, differential expression of specific markers of these putative malignant clusters was apparent exemplified by *PTCH1, MEIS1, POU3F4, FGFBP3, TWIST1* and *LHX9* that displayed low or no expression in non-malignant cells in co-culture and in the control organoid (Fig. 2B, D and Supplementary Table 6). Noting that *POU3F4* and *FGFBP3* have been linked to a neural stem cell (NSC) identity,^38,39^ we compared DAOY cells in co-culture with the DAOY tumour spheroid sample for expression of NSC markers (Supplementary Fig. S5A). Multiple NSC markers were upregulated in DAOY cells in co-culture including *SOX2*, *NES*, *FABP7* and *MSI1* (Fig. 2C). A similar trend was observed for ONS-76 cells in the three conditions, although the upregulation of these NSC markers in ONS-76 cells in co-culture was not as great (Fig. 2E).

**Figure 2.**
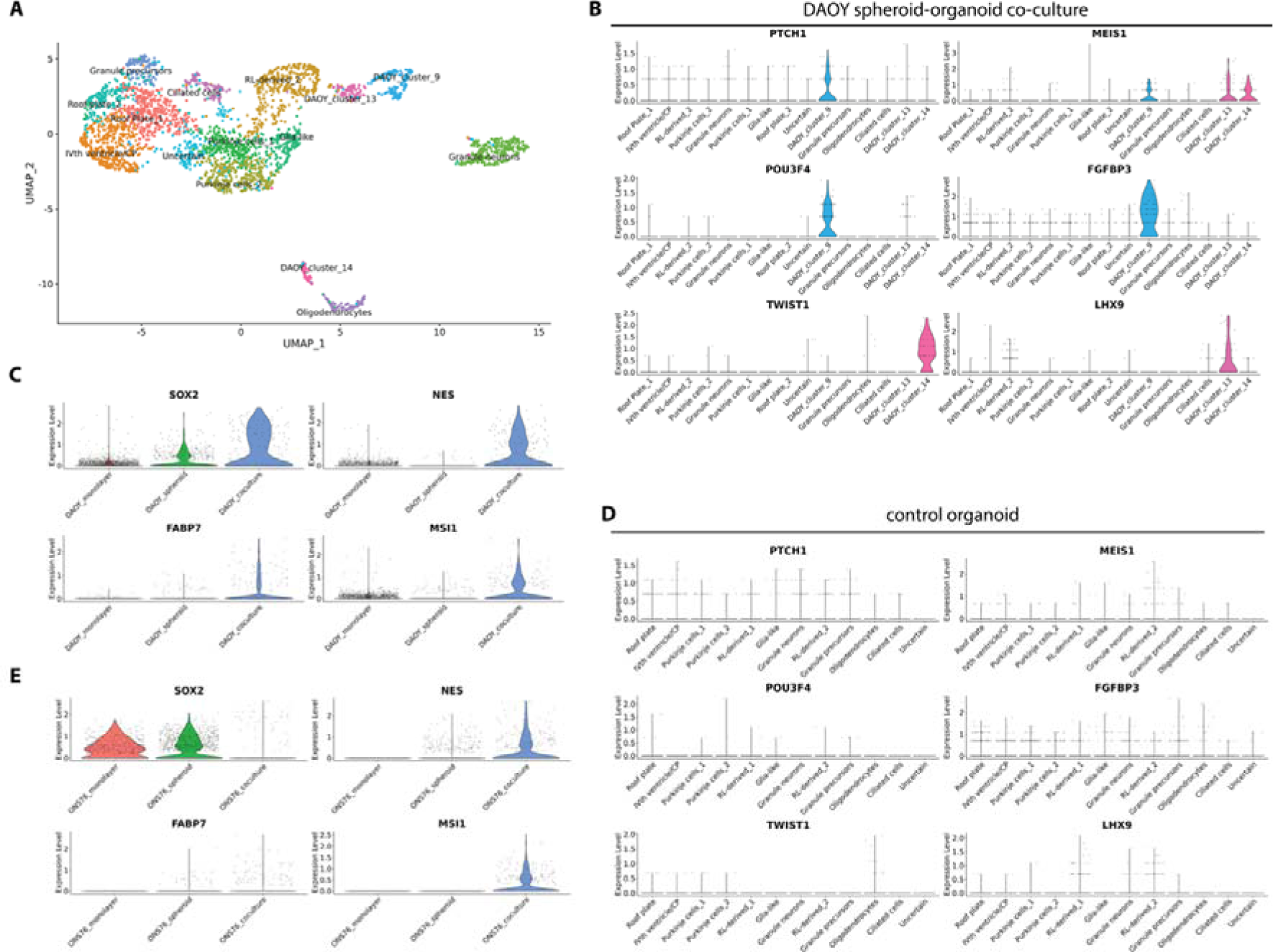
Upregulation of neural stem cell markers in DAOY malignant cells in co-culture. (A) UMAP visualisation of single-cell transcriptomes in DAOY tumour spheroid-organoid co-cultures shows three malignant clusters annotated as ‘DAOY_cluster_9, ‘DAOY_cluster_13’ and ‘DAOY_cluster_14’. (B) Violin plots showing expression of unique markers of malignant DAOY clusters in co-culture. (C) Violin plots of expression of the indicated neural stem cell markers in DAOY cells in tumour spheroid-organoid co-culture, tumour spheroid and monolayer cultures. (D) Expression of the indicated genes in cells of the control organoid sample. (E) Violin plots of expression of the indicated neural stem cell markers in ONS-76 cells in tumour spheroid-organoid co-culture, tumour spheroid and monolayer cultures.

The upregulation of NSC markers in malignant cells in our co-culture model is reminiscent of mouse models of SHH-MB, in which persistent SOX2-expressing granule cells designated as cancer stem cells (CSC) drive tumour formation and relapse.^6,40^ We therefore focused our analysis on further characterising the transcriptional profile of these cells. More granular analysis at the level of individual malignant clusters revealed striking differences in the expression of factors known to regulate *SOX2* compared to DAOY cells cultured in other conditions and ONS-76 cells (Fig. 3A).^41^ Only a single DAOY cluster in co-culture, ‘DAOY_cluster_9’ expressed multiple members of the *SOX2* regulatory network consisting of *POU3F2*, *ZIC2*, *OTX2*, *GLI2* and *SOX2* itself (Fig. 3B). This combinatorial pattern of gene expression was not observed in ONS-76 cells in co-culture (Fig. 3C), nor in tumour spheroid or monolayer samples of either cell line (Fig. 3D), or in non-malignant organoid cells (Fig. 3E). The co-expression of these markers in subpopulations of DAOY cells in co-culture suggested these cells adopted a new cell state in the presence of a non-malignant microenvironment.

**Figure 3.**
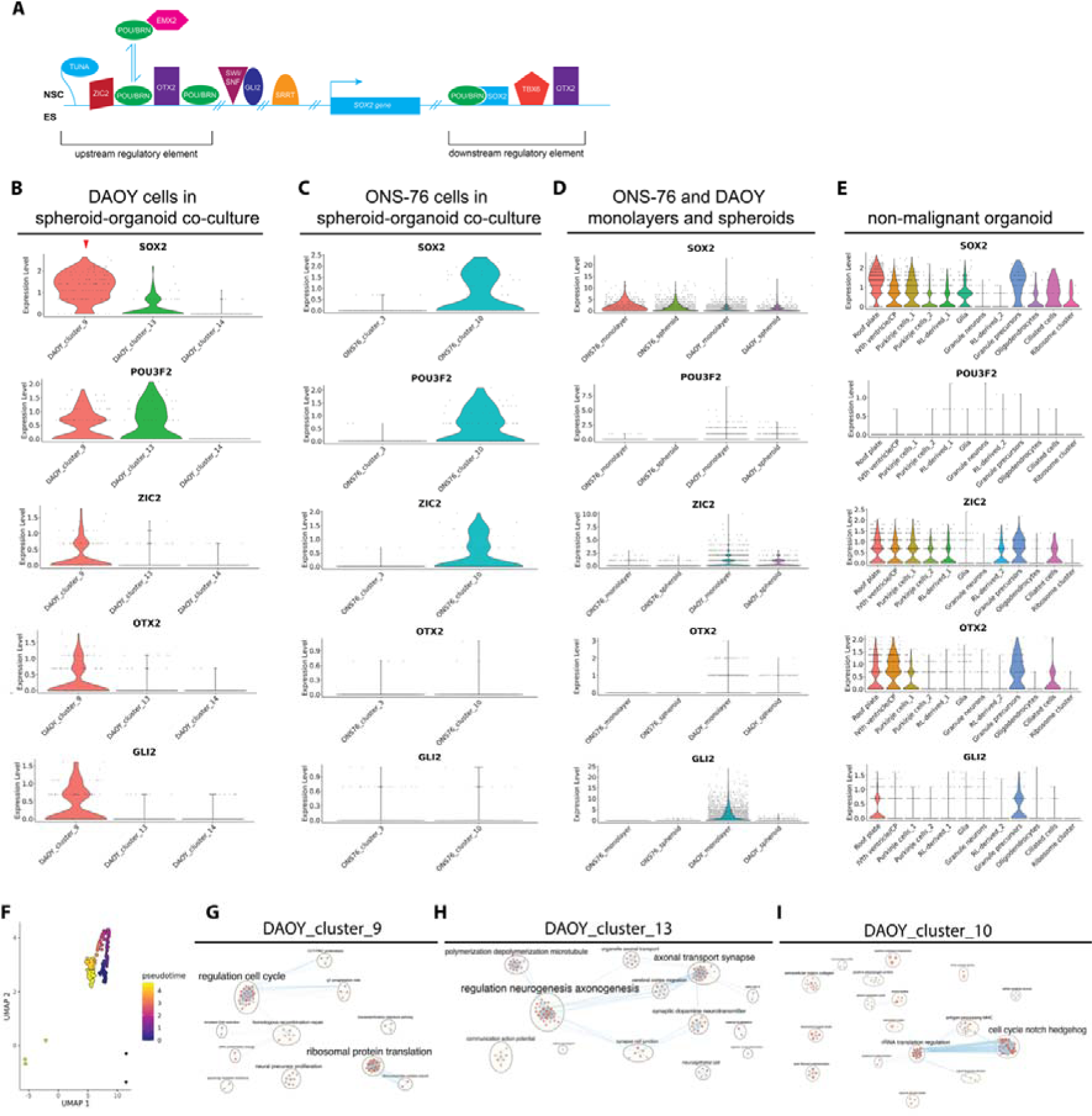
Multiple factors known to regulate the expression of *SOX2* in NSC and ES cells (ESC) are also expressed in a sub-population of DAOY cells in co-culture. (A) Schematic of the *SOX2* regulatory network in NSC and ESC (adapted from Bertolini et al.^41^). (B) Violin plots of expression of the *SOX2* regulatory network in ‘DAOY_cluster_9’ cells. (C) Expression of a sparse *SOX2* regulatory network in ONS-76 cells in tumour spheroid-organoid co-culture. (D) *SOX2* regulatory network expression in DAOY and ONS-76 monolayer and tumour spheroid cultures. (E) Gene expression of the *SOX2* regulatory network in non-malignant organoids. (F) Pseudotime trajectory of cells in ‘DAOY_cluster_9’. (G) Enrichment of GO biological pathway terms upregulated in ‘DAOY_cluster_9’ visualised in Cytoscape.^18^ (H) Enrichment of GO biological pathway terms upregulated in ‘DAOY_cluster_13’ visualised in Cytoscape. (I) Enrichment of GO biological pathway terms upregulated in ‘DAOY_cluster_10’ visualised in Cytoscape. *TBX6* was not expressed in malignant cells.

We hypothesised there could be an ontological relationship between the distinct DAOY cell states identified in co-culture. In particular, the UMAP plot of DAOY cells in co-culture (Fig. 2A) suggested ‘DAOY_cluster_9’ and ‘DAOY_cluster_13’ cells occupied opposite ends of a single trajectory. A pseudotime analysis supported the notion that ‘DAOY_cluster_9’ could undergo a progressive transition to a ‘DAOY_cluster_13’ cell state (Fig. 3F). Pathway enrichment analysis was consistent with the pseudotime analysis and showed that ‘DAOY_cluster_9’ cells were enriched for GO terms associated with a proliferative state (Fig. 3G; Supplementary Table 7), whereas ‘DAOY_cluster_13’ was enriched for neuronal differentiation terms (Fig. 3H and Supplementary Table 7). The last cluster, ‘DAOY_cluster_14’ was transcriptionally less similar to the other DAOY clusters (Fig. 2A). Enriched pathways for this cluster related to Notch and Hedgehog signalling, translation regulation and included extracellular matrix composition and wound healing gene sets in common with tumour cells cultured in isolation (Fig. 3I; Supplementary Table 7).

### CSC-like cells with a molecular profile resembling DAOY tumour cells in co-culture are present in patient SHH-MB

We analysed the expression of the *SOX2* regulatory network in a public scRNA-seq dataset of patient SHH-MB.^32^ Upon clustering tumour cells in Seurat ^13^ a single cluster in one of three SHH-MB tumours expressed a highly similar network in which *OTX2* was replaced by expression of another member of the regulatory network, *SRRT* (Fig. 3A and Fig. 4A).^41^ Furthermore, in one of two PDX models of SHH-MB from the same dataset,^32^ a single cluster expressed a common network, consisting of *POU3F2*, *OTX2*, *SRRT*, *GLI2* and *SOX2* (Fig. 4A). Given these differences, and to determine the prevalence and identity of cells expressing a signature of a *SOX2* regulatory network, we analysed a larger dataset of patient SHH-MB.^33^ Initial analysis of the data showed upregulation of all the aforementioned genes in malignant cells compared to the non-malignant microenvironment (Fig. 4B). By contrast, co-expression of these genes was weaker in other MB subtypes (Supplementary Fig. S5B). Clustering the individual SHH-MB tumours showed that in 7/8 samples two to four clusters emerged per tumour. One or more of these clusters expressed a *SOX2* regulatory network that was highly similar to the regulatory network in DAOY cells in co-culture (Fig. 4C, D; Supplementary Fig. S5C – H). The proportion of malignant cells expressing this signature ranged from 4.2% to 20.3% (median 8.8%) in this dataset.^33^ Quantification of the gene signature across MB subtypes showed enrichment in SHH-MB tumours (Fig. 4E). Furthermore, *TP53* mutation status (n = 2 *TP53* mutant SHH-MB tumours) was unrelated to the occurrence of the gene signature. These data confirmed that the patterns of gene expression in DAOY cells in co-culture were not simply an artifact related to our culture conditions, and instead suggested DAOY cells mirrored a CSC-like state found *in vivo*.

**Figure 4.**
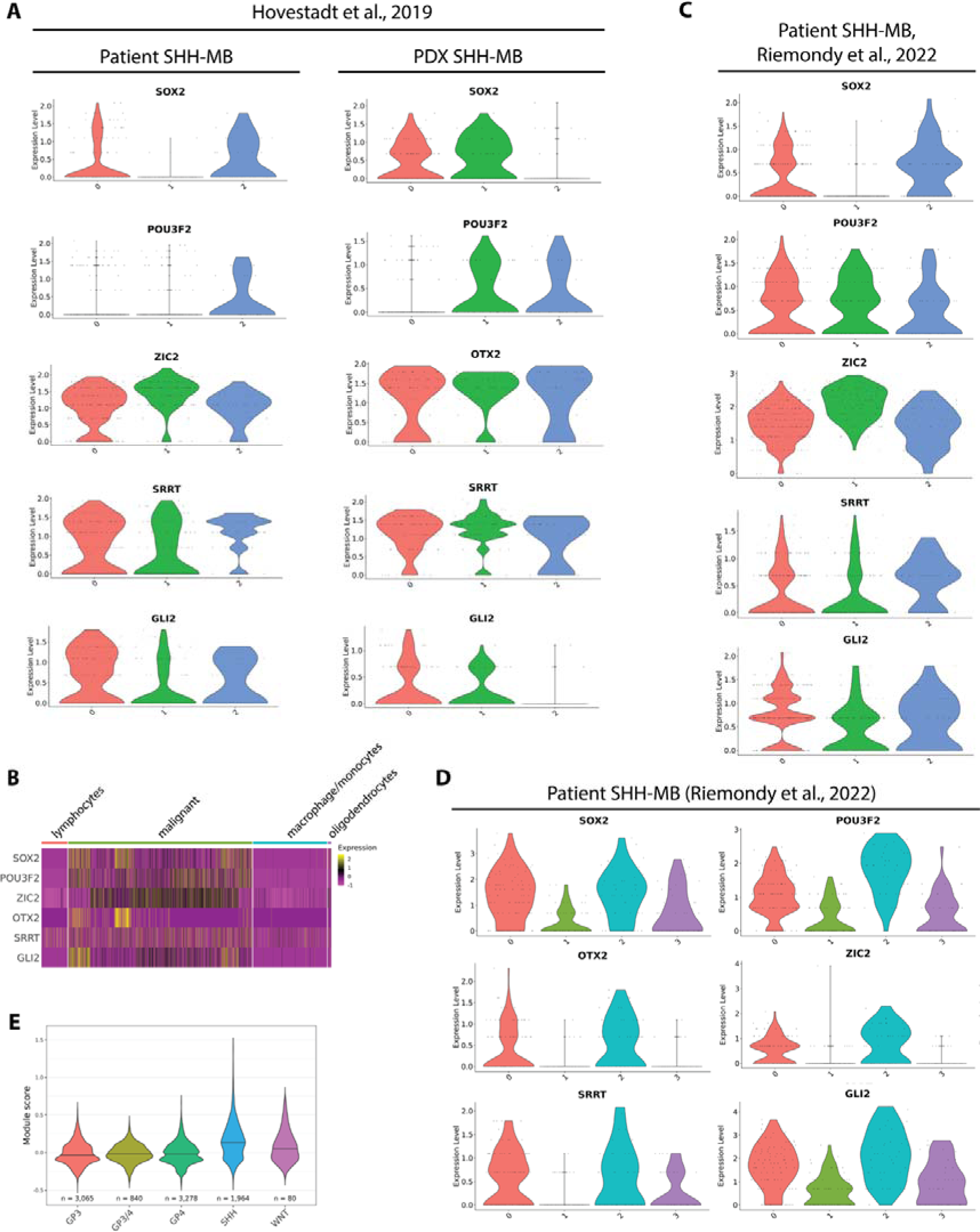
Expression of the *SOX2* regulatory network in patient SHH-MB datasets sequenced at single-cell level. (A) Violin plots of the expression of the *SOX2* regulatory network in one of three patient SHH-MB (tumour ID: SJ454) and in a SHH-MB PDX tumour model (tumour ID: RMCB18) from the same dataset.^32^ (B) Heatmap of expression of the *SOX2* regulatory network in malignant cells and in cells of the microenvironment in a second patient SHH-MB dataset.^33^ Cell labels are retained from the original publication. (C) Expression of the *SOX2* regulatory network in two of three UMAP clusters in a patient tumour (tumour ID: 1235) from the same dataset.^33^ (D) *SOX2* regulatory network expression in another tumour (tumour ID: 801) from the same dataset.^33^ (E) Violin plots of cell scores for the *SOX2* regulatory network in all medulloblastoma subtypes. Only malignant cells are included.^33^ For malignant cell numbers in each SHH-MB see Supplementary Fig. S4. *TBX6* was not expressed in malignant cells in either dataset.

### Greater transcriptional similarity of DAOY and ONS-76 cells in co-culture to patient SHH-MB

Encouraged by the transcriptional evidence suggesting tumour heterogeneity resembling SHH-MB *in vivo* in our organoid co-culture model, we assessed the transcriptional similarity of our model to patient tumours and PDX models. Therefore, we compared the computational integration of DAOY and ONS-76 tumour cells in co-culture and PDX models with patient SHH-MB tumours. We did not use canonical correlation implemented in Seurat for this part of the analysis because this algorithm excessively blends distinct cancer cell states.^21^ Instead, we employed the fastMNN algorithm,^42^ which performed well in an earlier benchmarking study of tumour datasets ^21^ to integrate three public SHH-MB patient single-cell datasets.^32^ Upon projection in UMAP space two major clusters emerged to which all three tumours contributed (Fig. 5A). Two SHH-MB PDX tumours ^32^ blended well with the patient tumours but showed greater mixing with one of the two major clusters, suggesting these PDX models did not recapitulate the full extent of transcriptional diversity of the patient tumours (Fig. 5B, C). Whereas DAOY and ONS-76 cells in co-culture, as expected, were less similar to patient tumours (Fig. 5D, E), in comparison with tumour spheroids of either cell line far greater similarity was evident (Fig. 5F, G). We also computed a ‘mixing metric’ to quantify how similar these datasets were after integration (Fig. 5H and Supplementary Table 8).^43^ Patient tumours scored the highest on this metric, followed closely by PDX tumours. Strikingly, our tumour spheroid organoid co-culture models also performed well, in contrast to tumour spheroid models.

**Figure 5.**
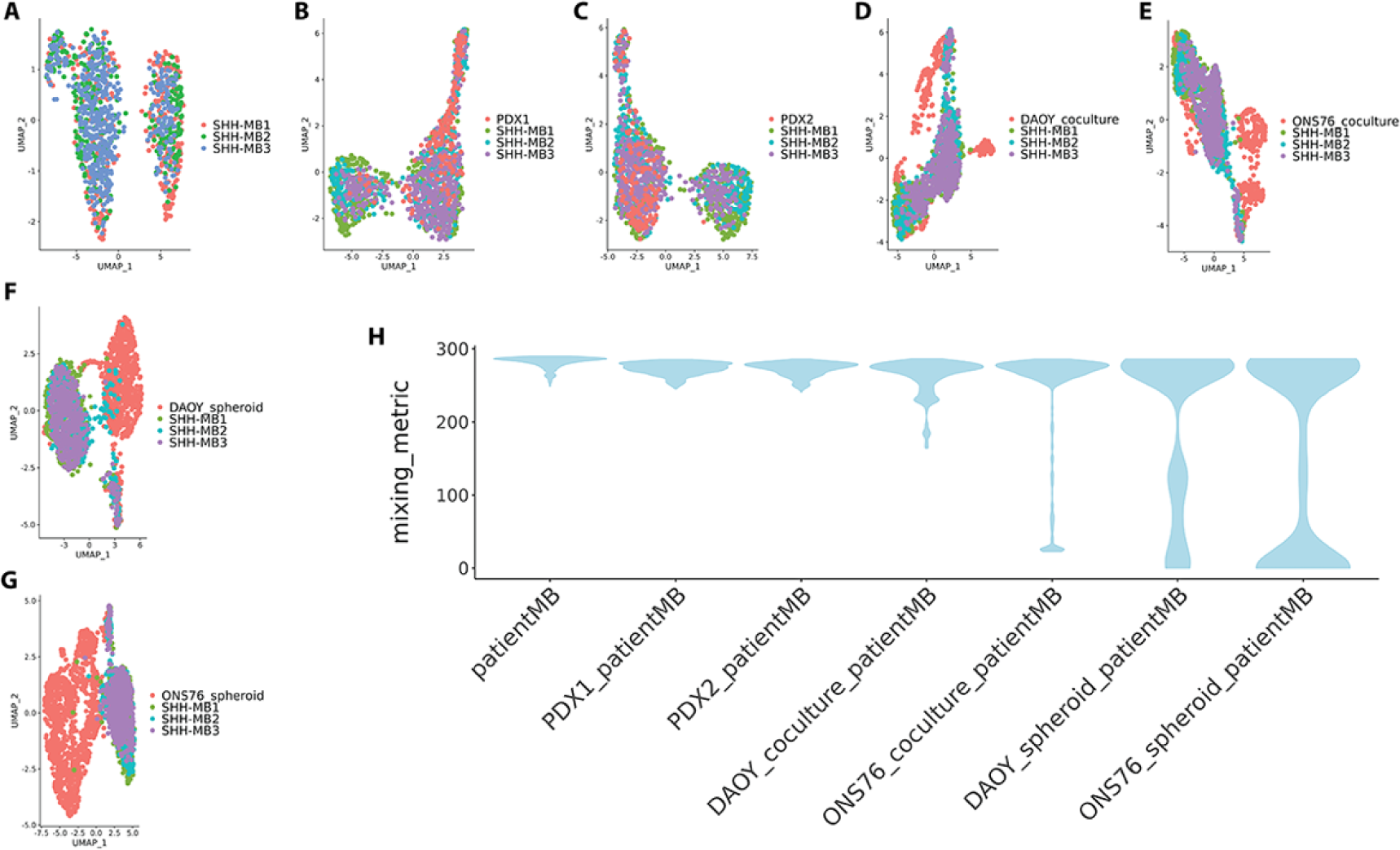
The co-culture tumour models recapitulate SHH-MB patient tumours better than conventional *in vitro* models. (A) UMAP visualisation of three SHH-MB patient tumours. (B) Integration and UMAP visualisation of cells from a PDX SHH-MB tumour (PDX1) with the same three SHH-MB patient tumours as in A. (C) Integration and UMAP visualisation of cells from a second PDX SHH-MB tumour (PDX2) with the three SHH-MB patient tumours. (D) Integration and UMAP visualisation of DAOY cells in tumour spheroid-organoid co-culture with the three SHH-MB patient tumours. (E) Integration and UMAP visualisation of ONS-76 cells in tumour spheroid-organoid co-culture with the three patient SHH-MB tumours. (F) Integration and UMAP visualisation of DAOY tumour spheroid cells with the three patient SHH-MB tumours. (G) Integration and UMAP visualisation of ONS-76 tumour spheroid cells with the three patient SHH-MB tumours. (H) Violin plots of mixing metric scores of the integrations shown in A to G. Patient and PDX tumour datasets were obtained from the published study by Hovestadt et al.^32^ Only malignant cells were used for scoring.

## Discussion

Human cerebellar organoid-based *in vitro* models have the potential to recapitulate patient MB more faithfully compared to conventional *in vitro* models.^44^ Here, we tested the importance of the non-malignant microenvironment in regulating malignant cell states by comparing the phenotypes of two long-established SHH-MB lines grown in co-culture or conventionally. Compared to conventional *in vitro* models, we detected greater maturity of malignant cells in co-culture. In particular, a subpopulation of ONS-76 cells activated the expression of *NEUROD1*, a canonical marker of granule cells, only upon co-culture. Additionally, a subpopulation of DAOY cells in co-culture expressed CSC-like markers and pseudotime and functional enrichment analyses were concordant attesting to their capacity for maturation into cells with neuronal features. The expression of CSC-like markers and upregulation of neuronal differentiation genes by malignant cells in co-culture revealed the emergence of cell states that were not observed in tumour cells cultured in isolation, under our conditions. Given that similar cell states are found in mouse models, PDX models and upon patient tumour molecular profiling,^6,32,40,45^ we further demonstrated the convergence of two SHH-MB cell lines towards *in vivo* cellular phenotypes. Therefore, the microenvironment can non-cell autonomously regulate SHH-MB cell states. Moreover, our computational analysis effectively addresses the challenge of the high non-malignant to malignant cell ratio in our co-cultures. The resilience of Seurat to cell downsampling made the identification of patient tumour-relevant cell states in our co-culture model possible.^12^

Given that malignant cell phenotypes in tumour spheroid-organoid co-culture were similar to those seen *in vivo* in patient tumours and PDX models (Fig. 1D, E; Fig. 3B; Fig. 4; Fig. 5) confirms the validity of our model in recapitulating patient tumours better than conventional *in vitro* models. Moreover, our findings suggest that the consensus view that these cell lines are poorly representative of patient tumours does not hold true when these malignant cell lines are cultured in a more physiological context. The emergence of new cell states in these long-established SHH-MB cell lines that was not evident in monolayer or tumour spheroid culture conditions was striking, more so given the relatively short duration (25 days) of co-culture. The immediate cause of these rapid phenotypic changes is likely to be transcriptional plasticity, reflecting functional intra-tumour heterogeneity that is shared by diverse malignant tumours.^46^

Malignant granule cells can differentiate in patient SHH-MB and in mouse models of this disease.^32,33,45,47,48^ Interestingly, the activation of *NEUROD1* in SHH-MB patient samples showed no dependency on *TP53* mutational status (Supplementary Fig. S4D), suggesting that differentiation processes can still be initiated in the presence of somatic *TP53* mutations.^49^ *NEUROD1* was upregulated in a sub-population of ONS-76 cells in tumour spheroids in co-culture with organoids compared to tumour monocultures (Fig. 1D, E). In both healthy and malignant cells, expression of *NEUROD1* was associated with actively cycling cells and cells with neuronal features (Fig. 1G, I; Supplementary Fig. S3C – F). Interestingly, however, in the malignant sub-population, upregulation of *NEUROD1* was not as tightly coupled with neuronal differentiation compared to non-malignant granule cells. These differences could reflect divergence of the properties of ONS-76 cells over an extended period *in vitro*, favouring the survival of cycling cells over more differentiated cells. Recent analysis of human granule cell ontogeny has demonstrated that, in contrast to mouse, these cells express *NEUROD1* during their cycling phase.^31^ The expression of this gene in a smaller proportion of non-malignant cycling granule precursors is consistent with the latter finding and further validates our cerebellar organoid model.

Our analysis showed that the DAOY CSC-like cells present in co-culture were enriched for pathways related to the cell cycle, which suggested they were proliferative (Fig. 3G and Supplementary Fig. S3C). Moreover, closely similar or equivalent cell states as defined by expression of the *SOX2* regulatory network were present in a majority of samples across two published scRNA-seq datasets that spanned infant to adult subtypes (Fig. 4 and Supplementary Fig. S5C – H).^32,33^ Reasons for the differences in the expressed members of the *SOX2* regulatory network in CSC-like subpopulations are unclear. Differences could relate to redundancy of distinct *SOX2* regulators or reflect divergence of functional states within CSC-like populations, associated with, for example, different proliferation rates and sensitivity to therapy. The traditional view is that CSC are rare, slow-cycling cells that are relatively quiescent compared to other malignant cells, and such tumour-initiating cells were identified in a SHH-MB mouse model.^6,40^ The proportion of CSC-like cells in the scRNA-seq dataset we analysed ^33^ was up to ∼20% with a median close to 10%, suggesting these cells might be more common than previously thought. Moreover, in another brain tumour, glioma, stem cells have also been reported to be proliferative, supporting the view that CSC states are non-uniform.^4^ Future functional studies are needed to determine if the SHH-MB cells with the signature we identified meet the standard for stemness in *in vivo* limiting dilution assays. Pseudotime and functional enrichment analyses were consistent and suggested these cells underwent a maturation process, associated with a continuum of cell states that led to the acquisition of neuron-like features (Fig. 3F – H). Furthermore, we discounted the possibility that the emergence of a CSC-like state in DAOY cells was caused by the *TP53* mutation because of the prevalence of a CSC-like state in patient SHH-MB analysed here that were wild type for *TP53* and the expression of NSC markers in ONS-76 cells in tumour spheroid-organoid co-culture (Fig. 2E).

In our co-culture model, single or multiple constituents of the non-malignant microenvironment could induce the emergent phenotypic states of SHH-MB cells. Regulatory interactions between malignant and non-malignant components that exploit local signalling networks are a well-established mechanism for tumour cells to gain a growth advantage.^50^ Microenvironmental factors implicated in these signalling networks *in vivo* include stromal cells, blood and lymphatic vessels, immune-inflammatory cells and extracellular matrix (ECM). Of these, ECM is the only one of these factors represented in our co-culture model and has a well-recognised role in tumour progression.^51^ Whether other components of the microenvironment in our model, namely progenitors, neurons, glia, roof plate or choroid plexus can produce malignant cell state-reprogramming signals is unclear. Identification of this factor(s) in future could open new therapeutic options.

## Authorship

JJ conceived the project and JJ and MvE designed experiments. MvE performed the experimental work and AN and JJ performed data analysis. EB and BSB provided advice on experimental design and analysis. JJ wrote the manuscript with contributions from all the authors.

## Data Availability

scRNA-seq data have been deposited in the National Centre for Biotechnology Information Gene Expression Omnibus (GEO) database and are publicly accessible through GEO accession number GSE254917 (https://www.ncbi.nlm.nih.gov/geo/query/acc.cgi?acc=GSE254917). Analysis code is available at https://github.com/jjacob12/carp_analysis50K/tree/main/PrePrint.

## Acknowledgements

Computation used the Oxford Biomedical Research Computing (BMRC) facility, a joint development between the Wellcome Centre for Human Genetics and the Big Data Institute supported by Health Data Research UK and the NIHR Oxford Biomedical Research Centre. We thank Dr Jia-Ling Ruan (Department of Oncology, Medical Sciences Division, University of Oxford) for preparing SHH-MB cell lines for authentication.

