## Supplementary Information for "Sonic hedgehog medulloblastoma cells in co-culture with cerebellar organoids converge towards *in vivo* malignant cell states" (from another version of this preprint: version 1, 2024-04-02)

<u>**Supplementary Information: van Essen et al.**</u>

### Supplementary Figure 1

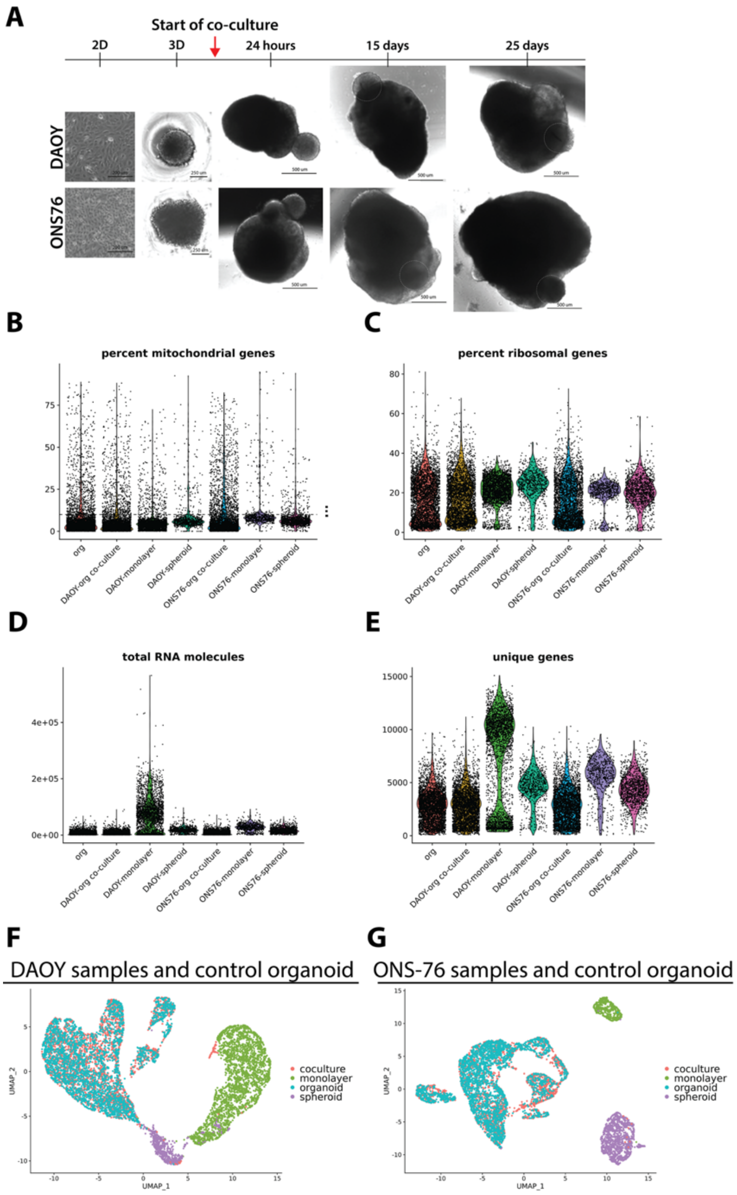


**Supplementary Figure 1**. QC metrics on samples used in this study. (A) Bright-field images of DAOY and ONS-76 monolayers, tumour spheroids and co-cultures with non-malignant cerebellar organoids. (B) Percent mitochondrial genes. (C) Percent ribosomal genes. (D) Total number of RNA molecules. (E) Number of unique genes. (F) UMAP visualisation of DAOY monolayer, spheroid, co-culture and control non-malignant cerebellar organoid samples. (G) UMAP visualisation of ONS-76 monolayer, spheroid, co-culture and control non-malignant cerebellar organoid samples.

**Supplementary Figure 2**

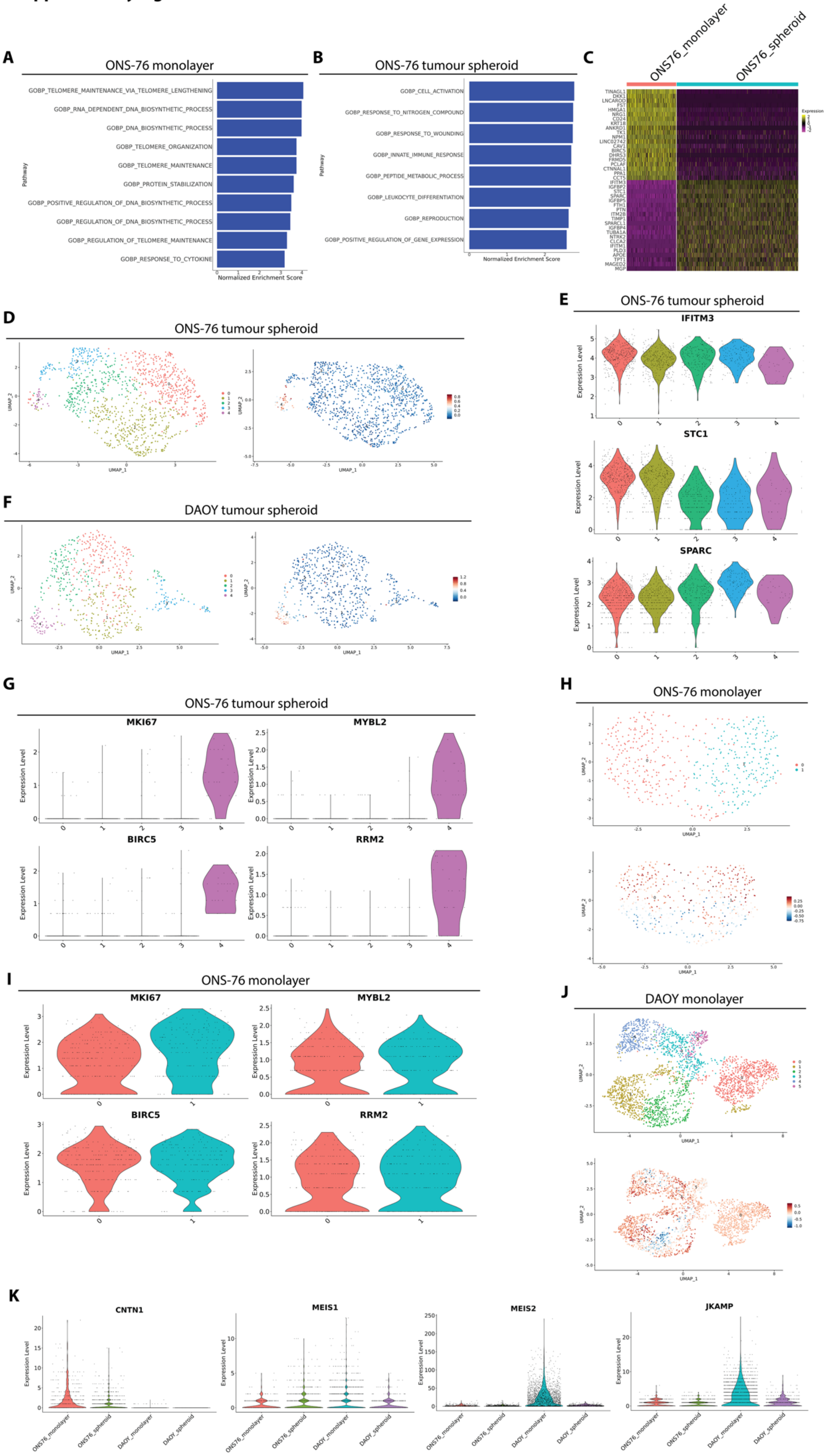

**Supplementary Figure 2**. Gene expression in ONS-76 and DAOY monolayers and tumour spheroids. (A) fgsea geneset enrichment of upregulated GO biological pathway terms evaluated by normalised enrichment scores in ONS-76 monolayer cells. (B) fgsea geneset enrichment of upregulated GO biological pathway terms evaluated by normalised enrichment scores in ONS-76 tumour spheroids. (C) Heatmap of top twenty differentially expressed genes in ONS-76 monolayer and tumour spheroid samples. (D) UMAP visualisation of ONS-76 tumour spheroid single cell transcriptomes identified five transcriptional cell clusters, 0 – 4 (left panel). Cell proliferation scores `are superimposed on the UMAP visualisations (right panel). (E) Violin plots of expression of the indicated genes in transcriptional clusters of ONS-76 spheroids that are associated with epithelial mesenchymal transition and stroma production. (F) UMAP visualisation of single cell transcriptomes of DAOY tumour spheroids identified five transcriptional cell clusters, 0 – 4 (left panel). Cell proliferation scores are superimposed on the UMAP visualisations (right panel). (G) Violin plots of the expression of the indicated genes in ONS-76 tumour spheroid clusters. (H) UMAP visualisation of single cell transcriptomes of ONS-76 monolayer cells identified two transcriptional clusters, 0 – 1 (upper panel). Cell proliferation scores are superimposed on the UMAP visualisations (lower panel). (I) Violin plots of gene expression in ONS-76 monolayer transcriptional clusters. (J) UMAP visualisation of single cell transcriptomes of ONS-76 monolayer cells identified six transcriptional clusters, 0 – 5 (upper panel). Cell proliferation scores are superimposed on the UMAP visualisations (lower panel). (K) Expression of differentiation markers in ONS-76 and DAOY monolayer and tumour spheroid samples.

**Supplementary Figure 3**

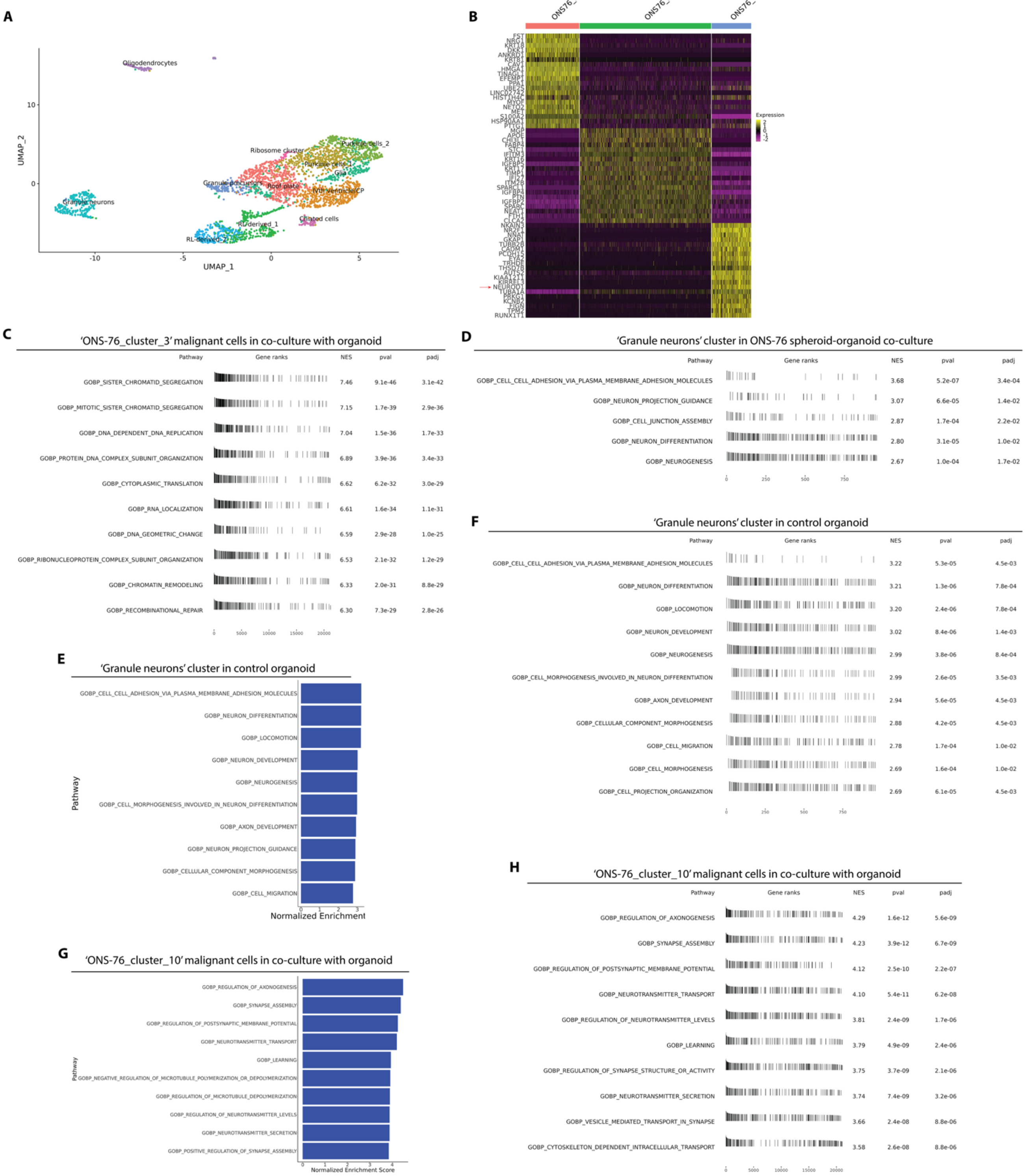


**Supplementary Figure 3**. Functional enrichment of GO biological pathways (GOBP) in ONS-76 malignant cells in tumour spheroid-organoid co-culture. (A) UMAP visualisation of single cell transcriptomes in non-malignant control organoid. (B) Heatmap of top twenty differentially expressed genes in ONS-76 monolayer, spheroid and malignant co-culture cells. Non-malignant cells in co-culture were not included. *NEUROD1* expression is indicated (red arrow). (C) GOBP enrichment scores and associated false discovery rate values (padj) in ‘ONS76_cluster_3’ cells in tumour spheroid-organoid co-culture. (D) GOBP enrichment scores and associated padj values in the ‘Granule neurons’ cluster in tumour spheroid-organoid co-culture. (E) fgsea geneset enrichment of upregulated GO biological pathway terms evaluated by normalised enrichment scores in the ‘Granule neurons’ cluster of the control organoid. (F) GOBP enrichment scores and associated padj values in the ‘Granule neurons’ cluster of the control organoid. (G) fgsea geneset enrichment of upregulated GO biological pathway terms evaluated by normalised enrichment scores in ‘ONS_cluster_10’ of the tumour spheroid-organoid co-culture. (H) GOBP enrichment scores and associated padj values in ‘ONS76_cluster_10’ cells in tumour spheroid-organoid co-culture.

**Supplementary Figure 4**

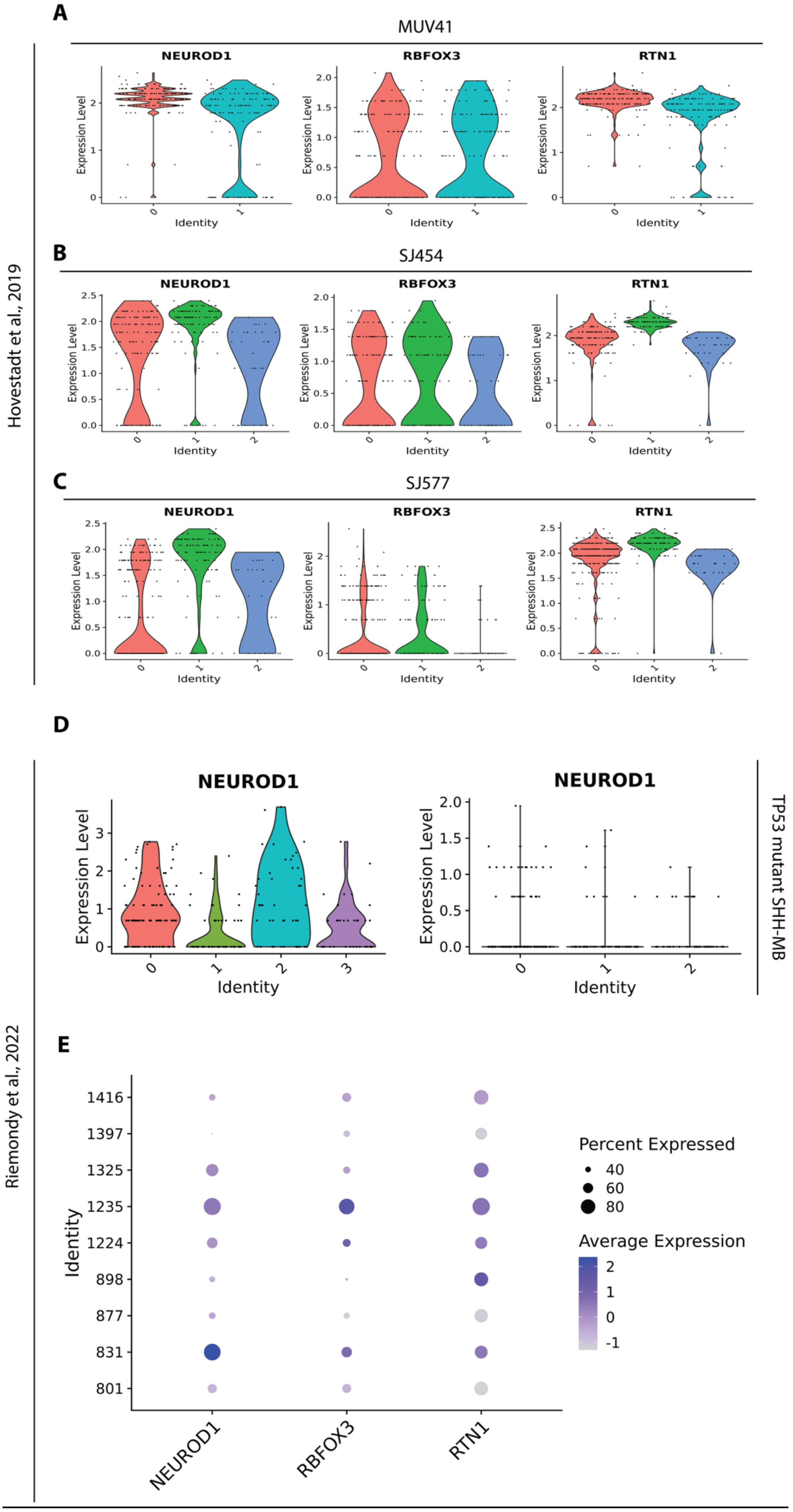

**Supplementary Figure 4**. Differentiating granule cell markers in SHH-MB tumours sequenced at single cell level across two public datasets.[1,2] (A) Violin plot of expression of the indicated markers in tumour sample MUV41 (n = 338 tumour cells).[1] (B) Violin plot of expression of the indicated markers in tumour sample SJ454 (n = 293 tumour cells).[1] (C) Violin plot of expression of the indicated markers in tumour sample SJ577 (n = 493 tumour cells).[1] (D) Violin plot of expression of *NEUROD1* in two *TP53* mutant tumours.[2] Left panel tumour ID: 801, right panel tumour ID: 1397. (E) Dot plot of the indicated granule cell differentiation markers expressed in nine SHH-medulloblastoma tumours (represented by identities 801 – 1416 on the y-axis).[2]

**Supplementary Figure 5**

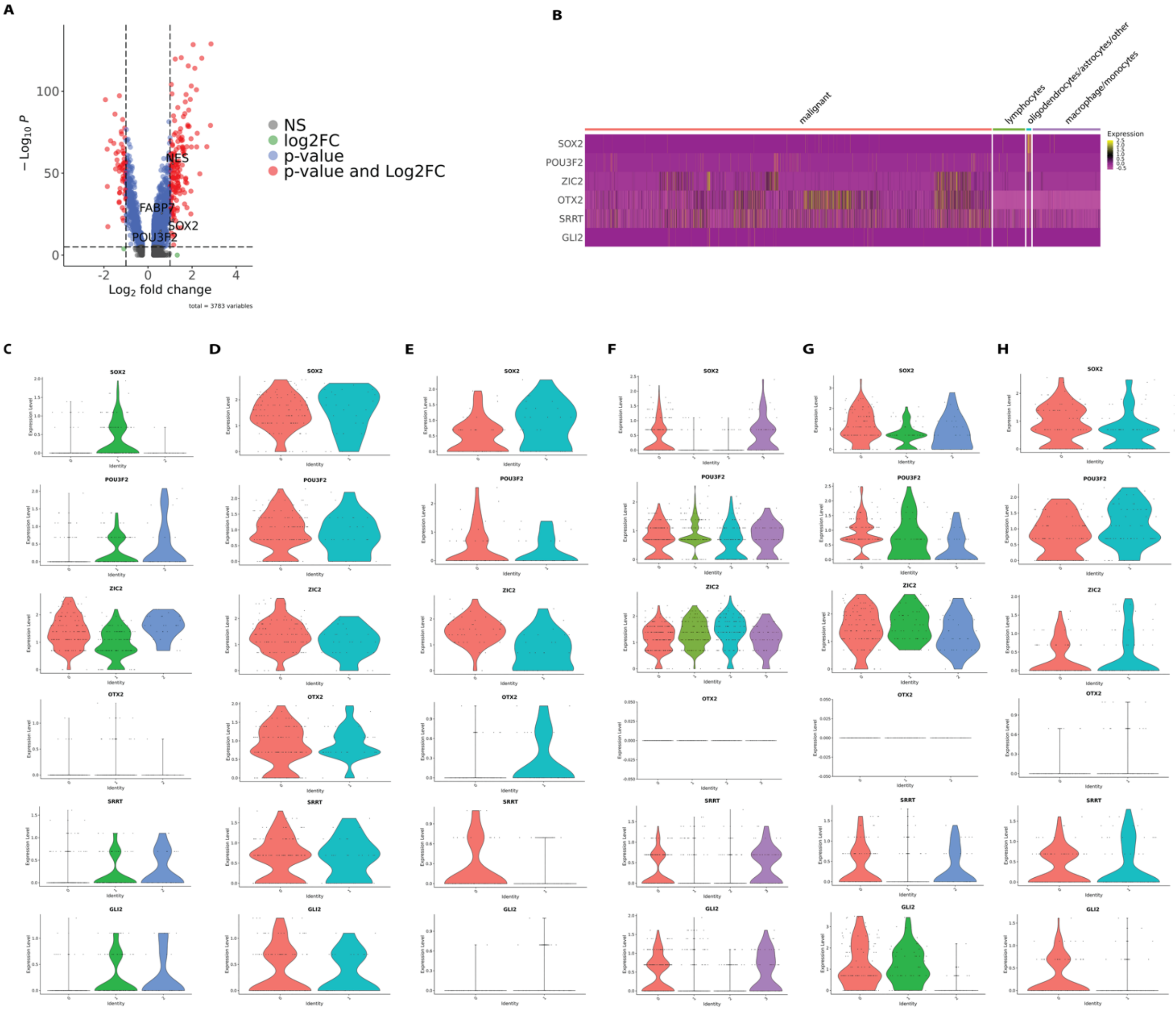

**Supplementary Figure 5**. Expression of stemness markers and a *SOX2* regulatory network in *in vitro* cultures and patient MB tumours. (A) Volcano plot of differentially expressed genes in DAOY cells in tumour spheroid-organoid co-culture compared to DAOY tumour-spheroids. NSC marker genes are upregulated. (B) Heatmap of the expression in single cells of the *SOX2* regulatory network for Group 3, Group 4 and WNT malignant and non-malignant cells. Cell labels are retained from the original publication. (C – H) Expression of the *SOX2* regulatory network in transcriptional cell clusters from a second dataset.[2] C, SHH-MB tumour ID: 831 (n = 264 tumour cells); D, SHH-MB tumour ID: 877 (n = 264 tumour cells); E, SHH-MB tumour ID: 898 (n = 66 tumour cells); F, SHH-MB tumour ID: 1325 (n = 538 tumour cells); G, SHH-MB tumour ID: 1397 (n = 198 tumour cells, *TP53* mutant); H, SHH-MB tumour ID: 1416 (n = 144 tumour cells).

**Supplementary Tables**

Supplementary Table 1. Authentication of DAOY-GFP and ONS-76-Luc-GFP cell lines by STR profiling.

Supplementary Table 2. Sample cell numbers.

Supplementary Table 3. Genes expressed in ONS-76 monolayer and ONS-76 tumour spheroid cells.

Supplementary Table 4. Gene set Enrichment for GO Biological Process terms in ONS-76 tumour spheroid cluster 0, ONS-76 tumour spheroid cluster 4 and ONS-76 monolayer cluster 0.

Supplementary Table 5. Gene set enrichment for GO biological process terms and for C6 oncogenic signature gene sets in integrated DAOY monolayer and tumour spheroid samples, and in individual UMAP clusters of DAOY monolayer and tumour spheroid samples.

Supplementary Table 6. Markers of ONS-76 and DAOY malignant cell clusters in tumour spheroid-organoid co-culture.

Supplementary Table 7. g:Profiler enriched pathways in DAOY cells in tumour spheroid-organoid co-culture.

Supplementary Table 8. Mixing metric summary statistics quantifying the extent of intermixing of patient SHH-medulloblastoma samples with other samples.
