## Supplementary figures and images for "Sonic hedgehog medulloblastoma cells in co-culture with cerebellar organoids converge towards *in vivo* malignant cell states" (from another version of this preprint: version 1, 2024-04-02)

### Supplementary table 1

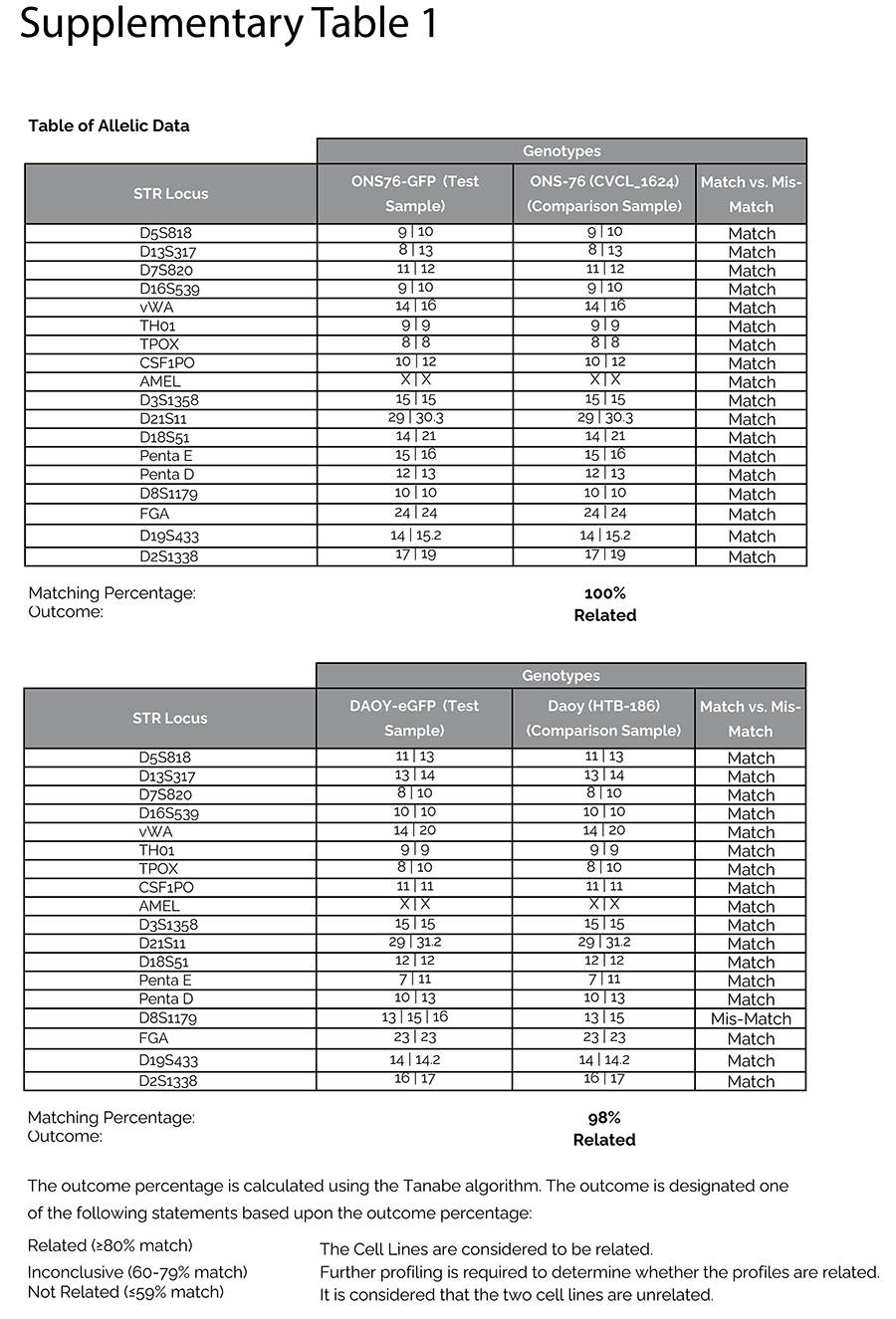
